# Saccharide Analysis of Onion Outer Epidermal Walls

**DOI:** 10.1101/2021.01.25.428129

**Authors:** Liza A. Wilson, Fabien Deligey, Tuo Wang, Daniel J. Cosgrove

## Abstract

**Background:** Epidermal cell walls have special structural and biological roles in the life of the plant. Typically they are multi-ply structures encrusted with waxes and cutin which protect the plant from dehydration and pathogen attack. These characteristics may also reduce chemical and enzymatic deconstruction of the wall for sugar analysis and conversion to biofuels. We have assessed the saccharide composition of the outer epidermal wall of onion scales with different analytical methods. This wall is a particularly useful model for cell wall imaging and mechanics.

**Results:** Epidermal walls were depolymerized by acidic methanolysis combined with 2 M trifluoracetic acid hydrolysis and the resultant sugars were analyzed by high-performance anion-exchange chromatography with pulsed amperometric detection (HPAEC-PAD). Total sugar yields based on wall dry weight were low (53%). Removal of waxes with chloroform increased the sugar yields to 73% and enzymatic digestion did not improve these yields. Analysis by gas chromatography/mass spectrometry (GC/MS) of per-*O*-trimethylsilyl (TMS) derivatives of the sugar methyl glycosides produced by acidic methanolysis gave a high yield for galacturonic acid (GalA) but glucose (Glc) was severely reduced. In a complementary fashion, GC/MS analysis of methyl alditols produced by permethylation gave substantial yields for glucose and other neutral sugars, but GalA was severely reduced. Analysis of the walls by ^13^C solid-state NMR confirmed and extended these results and revealed 15% lipid content after chloroform extraction (potentially cutin and unextractable waxes).

**Conclusions:** Although exact values vary with the analytical method, our best estimate is that polysaccharide in the outer epidermal wall of onion scales is comprised of homogalacturonan (~50%), cellulose (~20%), galactan (~10%), xyloglucan (~10%) and smaller amounts of other polysaccharides. Low yields of specific monosaccharides by some methods may be exaggerated in epidermal walls impregnated with waxes and cutin and call for cautious interpretation of the results.

## Background

The outer epidermal wall of stems and leaves has distinctive physical and structural properties connected with its role to limit growth of these organs and to protect them (1–5). Typically the outer epidermal wall is a multi-lamellate structure impregnated with hydrophobic substances (cutin, waxes) and sometimes lignified (5–8). These structural features provide a measure of physical resistance to water loss and damage by pathogens and insects. Impregnation with hydrophobic substances may present special difficulties for wall saccharide analysis by interfering with solubilization and depolymerization of wall polysaccharides prior to monosaccharide analysis. Such impediments are analogous to obstacles encountered during cellulosic biomass conversion to biofuels.

This study concerns monosaccharide analysis of the outer periclinal wall of onion scale epidermis. This wall is ~7 μm thick (9) and comprised of many lamellae in which cellulose is organized in highly anisotropic 2D networks (10) that when averaged across the total thickness of the wall can appear to be isotropic in some cases (11) and slightly anisotropic in other cases (12, 13), depending on developmental state. Because of the simplicity of preparing epidermal strips, this material has frequently been studied for optical and mechanical analyses (14–17). When the abaxial (‘lower’) epidermis is peeled, a relatively clean strip can be obtained that consists predominantly of the outer periclinal wall (18, 19). This relatively homogeneous wall preparation has been used to assess the role of different wall components in wall mechanics by a combination of selective enzymatic digestions and imaging by atomic force microscopy (20). Although the monosaccharide composition of onion parenchyma walls has been well studied (21–26), there is less information about onion epidermal walls and none as far as we know specifically focused on the outer periclinal wall, which has been the subject of detailed mechanics, microscopy and spectroscopy studies recently (9, 11, 20, 27–33). In preliminary attempts to measure total sugars in this material by the phenol/sulfuric acid method (34), we discovered that a colored reaction product persistently interfered with this colorimetric method. Consequently we explored other methods for saccharide analysis of this material.

Here we report the results of these analyses, based on different ‘wet-bench’ approaches. We compared acid hydrolysis methods followed by HPAEC-PAD to assess monosaccharide compositions and tested whether removal of surface waxes by chloroform extraction would improve results. We also investigated whether treatment with a cell wall degrading enzyme gave better yields than chemical hydrolysis. These results were compared with two methods of GC/MS and with analysis by solid-state NMR. The latter method does not require solubilization or hydrolysis of the wall polysaccharides, which commonly entail a tradeoff between complete hydrolysis of the polysaccharides and chemical degradation of saccharides (35). Selective losses in these steps are commonly a problem in standard approaches to analysis of plant cell wall composition and are potentially aggravated in epidermal walls where impregnation with cutin and waxes and cellulose bundling may reduce the efficiency of wall depolymerization. This point resembles lignocellulose recalcitrance to chemical and enzymatic depolymerization that limits biofuel production (36–39)

## Results

Outer epidermal walls were peeled from the abaxial side of onion scales, washed with buffered detergent to remove adherent cytoplasmic debris, de-starched with amylase, and freeze dried prior to analysis (see Materials and Methods for details). Saccharide loss to the wash solution was negligible (total of 0.1% of the initial dry weight). Because the hydrated wall is thin (~7 μm), we deemed it to be readily accessible to external solutions and did not grind the epidermal peels; this avoided grinding losses that may be substantial and variable when small amounts of material are handled. Our first analytical attempts used chemical hydrolyses of epidermal walls followed by monosaccharide analysis by HPAEC-PAD (Figure 1). These results were compared with two methods based on GC/MS and with analysis by ^13^C solid-state NMR.

**Figure 1:**
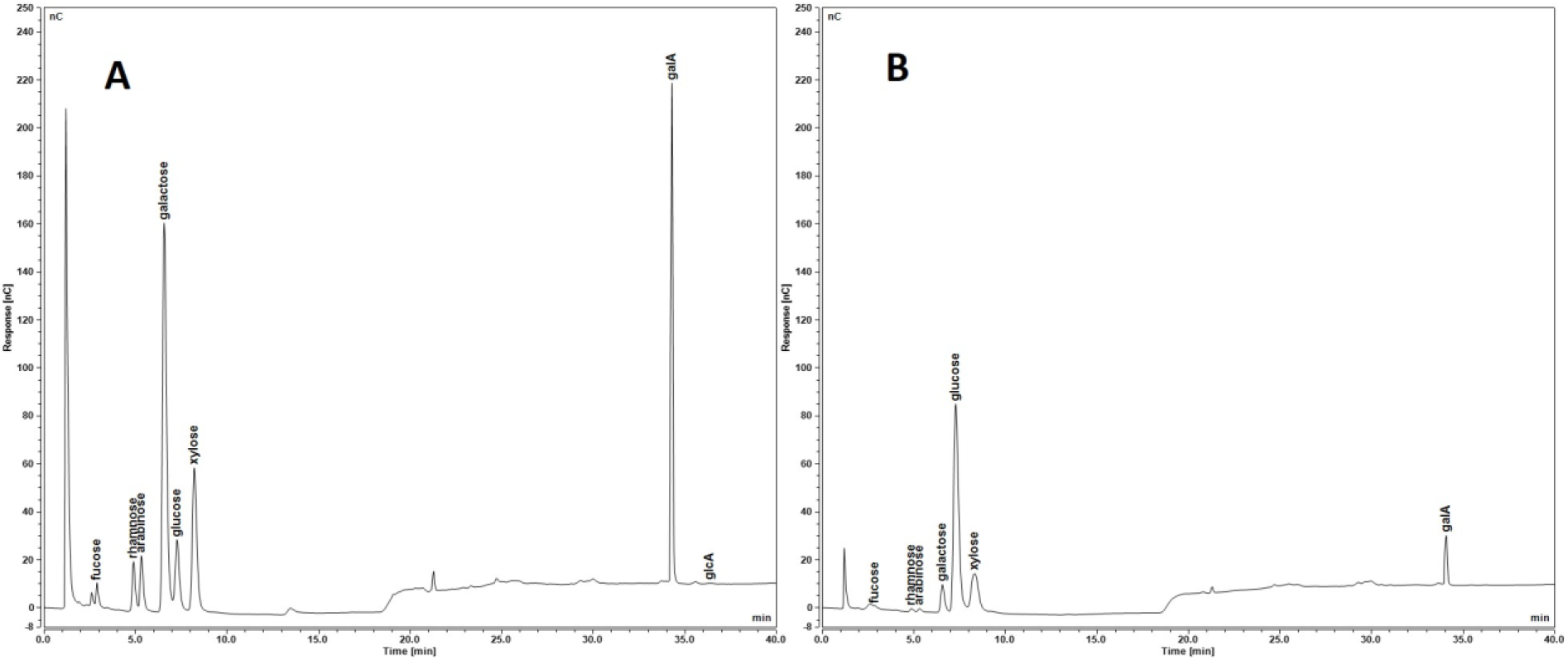
Representative HPAEC-PAD chromatograms from saccharide analysis of onion epidermal peels separated on a CarboPac PA20 analytical column with monosaccharides of interest labeled. **A**, soluble fraction after met/TFA hydrolysis; **B**, insoluble fraction of met/TFA hydrolysis further analyzed with 2-step sulfuric acid hydrolysis, neutralized, and separated on the column. GalA = galacturonic acid; GlcA = glucuronic acid.

### Chemical hydrolysis followed by HPAEC-PAD analysis

HPAEC-PAD is a sensitive and relatively simple method, requiring fewer steps than gas chromatography which entails chemical derivatization of the sugars (35, 40, 41). Efficient chemical hydrolysis of whole walls is challenging because of the heterogenous nature of the wall polysaccharides and their packing in insoluble forms within the wall. Harsh hydrolysis conditions may lead to saccharide degradation while mild conditions may lead to incomplete hydrolysis (42). To depolymerize non-crystalline polysaccharides, we used acidic methanolysis followed by hydrolysis in hot 2 M trifluoroacetic acid (met/TFA) (35). This tandem method reportedly improves hydrolysis of GalA-containing pectic polysaccharides without excessive degradation of neutral sugars. The cellulosic residue was then hydrolyzed with sulfuric acid in a two-step process (Saeman’s hydrolysis (43)).

As displayed in Table 1, GalA was the dominant monosaccharide detected in this analysis (51% of total saccharides), followed by Glc (24%), galactose (Gal, 13%) and smaller amounts of other monosaccharides. The HPAEC-PAD protocol does not separate xylose from mannose (Man), but GC/MS analysis (below) indicates this peak is 55% Xyl, 45% Man, so we estimate 4.3% Xyl and 2.1% Man in the epidermal walls. The results in this table account for an average of 532 μg of saccharide per mg of dry wall, suggesting the presence of other substances in the wall or larger yield losses than estimated.

**Table 1:** Saccharide composition of onion epidermal wall by met/TFA/Saeman’s hydrolysis.

| | | $\mu\text{g}/\text{mg}$ dry weight | | | | | | | | | |
| --- | --- | --- | --- | --- | --- | --- | --- | --- | --- | --- | --- |
|  |  | Trial 1 |  | Trial 2 |  | Trial 3 |  | mean |  | total | % |
|  |  | mTFA | Sae | mTFA | Sae | mTFA | Sae | mTFA | Sae |  |  |
| Glc |  | 6.9 | 21.6 | 13.1 | 144.9 | 17.1 | 186.3 | 12.3<br>(2.3) | 117.6<br>(19.8) | 130.0 | 24 |
| GalA |  | 183.5 | 16.5 | 255.5 | 8.5 | 321.3 | 29.0 | 253.4<br>(23.1) | 18.0<br>(2.4) | 271.5 | 51 |
| Gal |  | 64.4 | 1.9 | 52.0 | 2.8 | 87.0 | 4.5 | 67.8<br>(7.2) | 3.0<br>(0.6) | 70.9 | 13 |
| Xyl/Man* |  | 27.8 | 6.7 | 21.7 | 7.6 | 29.5 | 8.8 | 26.3<br>(1.8) | 7.7<br>(0.4) | 34.0 | 6 |
| Rha |  | 14.8 | 0.5 | 5.4 | 1.0 | 12.9 | 1.0 | 11.0<br>(1.9) | 0.8<br>(0.1) | 11.9 | 2 |
| Ara |  | 8.0 | 0.2 | 5.1 | 1.0 | 11.1 | 0.8 | 8.1<br>(1.2) | 0.6<br>(0.2) | 8.7 | 2 |
| Fuc |  | 3.1 | 0.3 | 2.4 | 1.5 | 4.1 | 1.6 | 3.2<br>(0.4) | 1.1<br>(0.4) | 4.4 | 1 |
| GlcA |  | 0.8 | 0.0 | 0.0 | 0.0 | 1.8 | 0.0 | 0.9<br>(0.3) | 0.0<br>(0.0) | 0.9 | 0.2 |
| **total** |  | **309.3** | **47.7** | **355.1** | **167.3** | **485.0** | **232.0** | **383.1** | **149.0** | **532.1** | **100** |

mTFA, peel hydrolyzed with methanolysis with TFA
Sae, 2-step sulfuric acid (Saeman's hydrolysis) on residue
Xyl/Man\*, HPAEC-PAD parameters did not separate xylose and mannose peaks
$\mu\text{g}/\text{mg}$ values were adjusted using correction factors listed in Methods
(SEM), standard error of mean of 3 biological replicates

Impregnation of the outer epidermal wall with cutin and waxes is a potential reason for low saccharide yields in the above analysis. These hydrophobic substances resist acids and might protect the wall polysaccharides from acid hydrolysis, and they might contribute substantially to the mass of the wall. To test these ideas, we extracted the walls with chloroform to solubilize waxes, resulting in a 13% decrease in dry weight of the wall (mean of two measurements).

Saccharide analysis of the chloroform wash detected galacturonic acid in an amount equivalent to 1% of the wall and only trace amounts of other sugars. We infer that the remaining 12% decrease in wall mass was due to wax removal. The extracted residue was then chemically depolymerized by sequential met/TFA and Saeman’s hydrolyses and analyzed by HPAEC-PAD. As displayed in Table 2, the total sugar yield increased to an average of 726 μg per mg of wall. The most notable difference from Table 1 is in the yield of Glc, which nearly doubled because of a doubling of the yield for Saeman’s hydrolysis (from a mean of 149 to 284 μg Glc per mg). This result confirms that pretreatment of the epidermal peel with an organic solvent, chloroform, can increase the hydrolyzability of the wall, especially in the cellulose-rich fraction. This suggests that much of the cellulose in the outer epidermal wall may be covered or embedded in waxes.

**Table 2:**
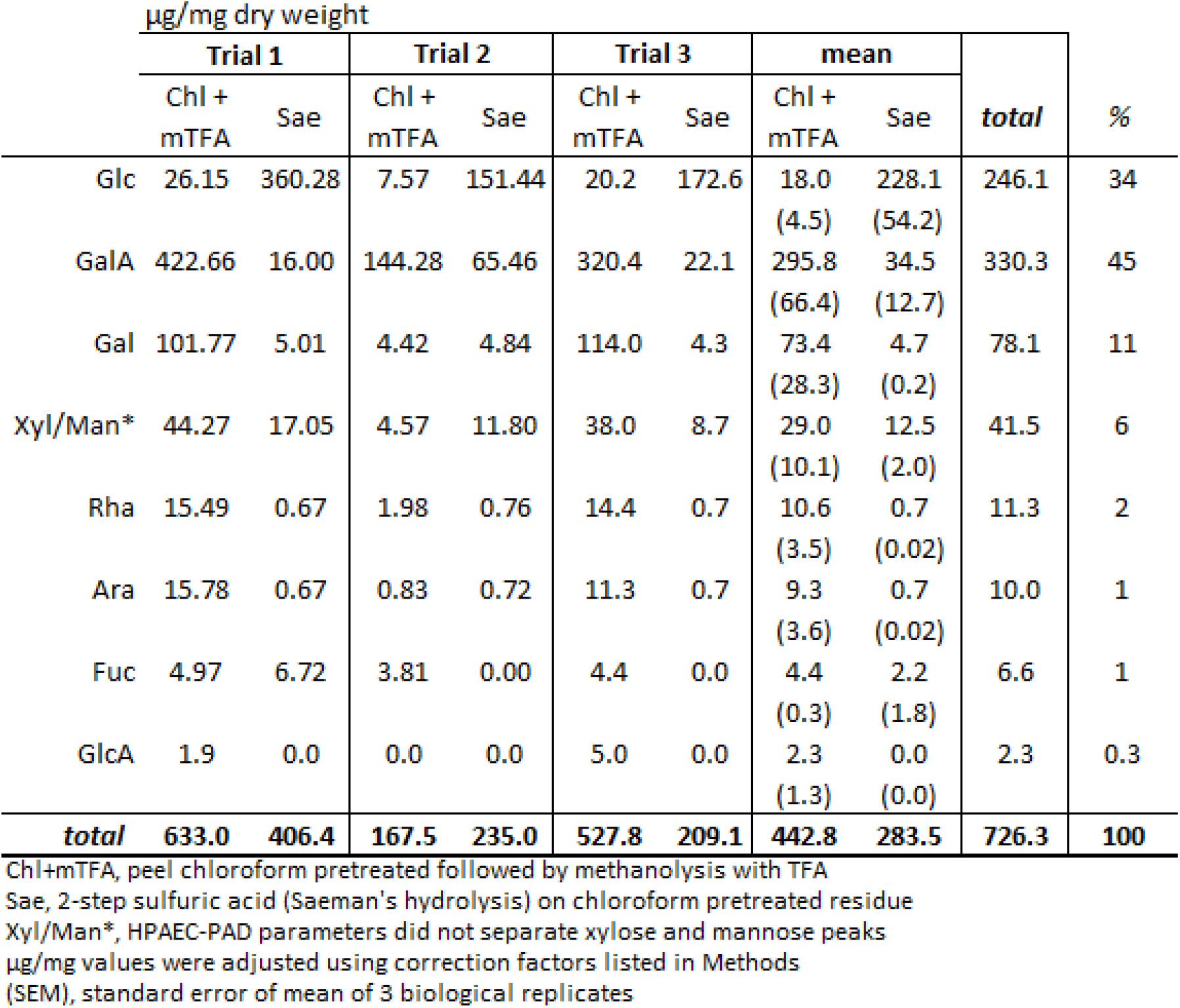
Saccharide composition of onion epidermal wall pre-extracted with chloroform and met/TFA/Saeman’s hydrolysis.

We next attempted to augment chemical hydrolysis of the wall by digestion with Driselase digestion prior to met/TFA and Saeman’s hydrolysis. Driselase is a complex mixture of wall hydrolases that degrades the wall to saccharide monomers and oligomers (44). As displayed in Table 3, Driselase pre-hydrolysis increased total sugar detected in met/TFA fraction, but at a cost to the Saeman’s fraction, with the result that the total sugar content was nearly the same as in Table 1 and less than in the dewaxed walls (Table 2). GalA and glucose along with galactose were again the predominant matrix saccharides.

**Table 3:**
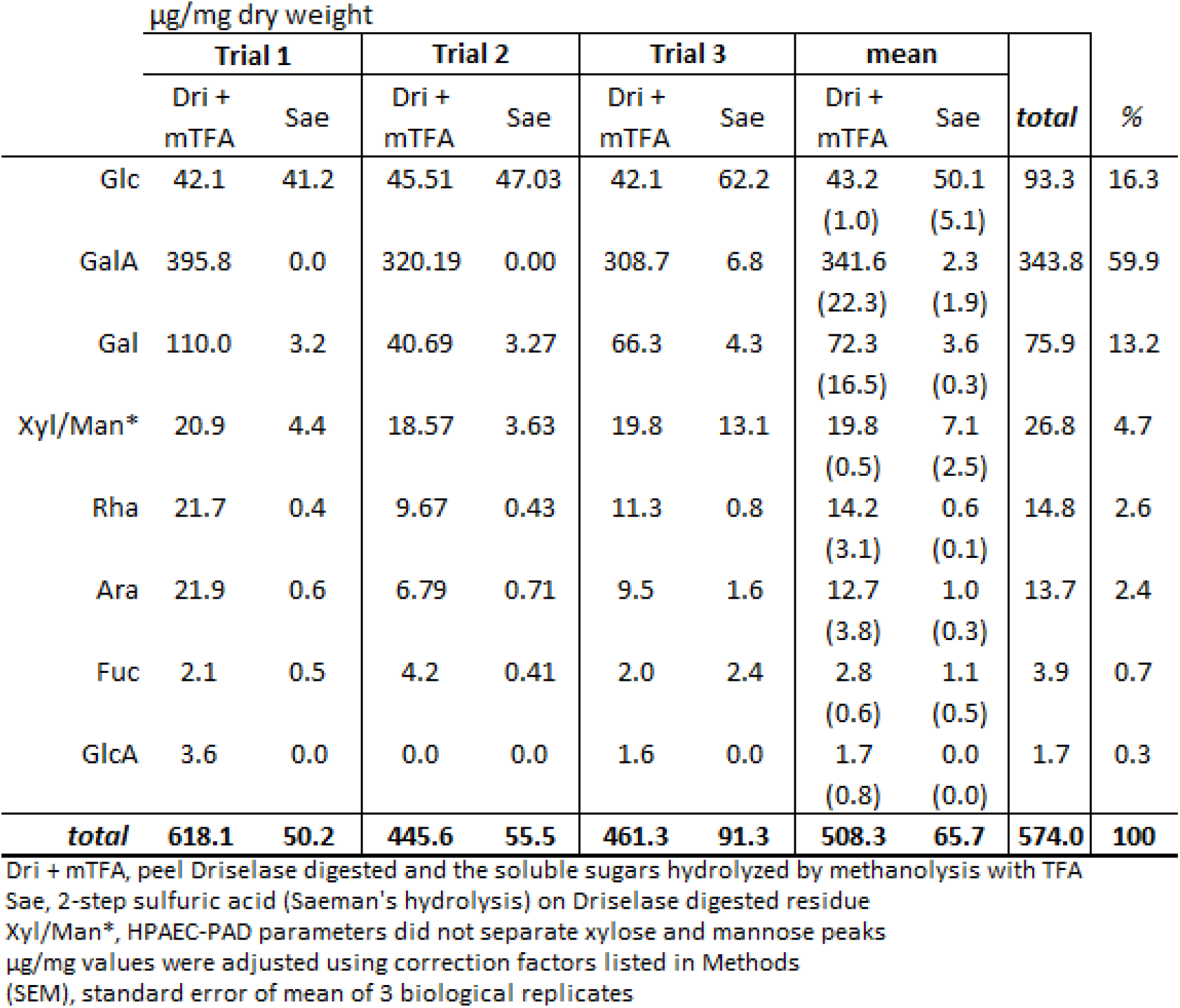
Saccharide composition of onion epidermal wall pre-digested with Driselase enzyme followed by met/TFA/Saeman’s hydrolysis.

| | | $\mu\text{g}/\text{mg}$ dry weight | | | | | | | | | |
| --- | --- | --- | --- | --- | --- | --- | --- | --- | --- | --- | --- |
|  |  | Trial 1 |  | Trial 2 |  | Trial 3 |  | mean |  | total | % |
|  |  | Dri + mTFA | Sae | Dri + mTFA | Sae | Dri + mTFA | Sae | Dri + mTFA | Sae |  |  |
| Glc |  | 42.1 | 41.2 | 45.51 | 47.03 | 42.1 | 62.2 | 43.2<br>(1.0) | 50.1<br>(5.1) | 93.3 | 16.3 |
| GalA |  | 395.8 | 0.0 | 320.19 | 0.00 | 308.7 | 6.8 | 341.6<br>(22.3) | 2.3<br>(1.9) | 343.8 | 59.9 |
| Gal |  | 110.0 | 3.2 | 40.69 | 3.27 | 66.3 | 4.3 | 72.3<br>(16.5) | 3.6<br>(0.3) | 75.9 | 13.2 |
| Xyl/Man* |  | 20.9 | 4.4 | 18.57 | 3.63 | 19.8 | 13.1 | 19.8<br>(0.5) | 7.1<br>(2.5) | 26.8 | 4.7 |
| Rha |  | 21.7 | 0.4 | 9.67 | 0.43 | 11.3 | 0.8 | 14.2<br>(3.1) | 0.6<br>(0.1) | 14.8 | 2.6 |
| Ara |  | 21.9 | 0.6 | 6.79 | 0.71 | 9.5 | 1.6 | 12.7<br>(3.8) | 1.0<br>(0.3) | 13.7 | 2.4 |
| Fuc |  | 2.1 | 0.5 | 4.2 | 0.41 | 2.0 | 2.4 | 2.8<br>(0.6) | 1.1<br>(0.5) | 3.9 | 0.7 |
| GlcA |  | 3.6 | 0.0 | 0.0 | 0.0 | 1.6 | 0.0 | 1.7<br>(0.8) | 0.0<br>(0.0) | 1.7 | 0.3 |
| **total** |  | **618.1** | **50.2** | **445.6** | **55.5** | **461.3** | **91.3** | **508.3** | **65.7** | **574.0** | **100** |

Dri + mTFA, peel Driselase digested and the soluble sugars hydrolyzed by methanolysis with TFA
Sae, 2-step sulfuric acid (Saeman's hydrolysis) on Driselase digested residue
Xyl/Man\*, HPAEC-PAD parameters did not separate xylose and mannose peaks
$\mu\text{g}/\text{mg}$ values were adjusted using correction factors listed in Methods
(SEM), standard error of mean of 3 biological replicates

We next combined the chloroform pre-extraction with the Driselase pre-digestion with the expectation of increasing sugar yields further. The results, shown in Table 4, proved counter to this expectation. As before, Driselase pre-digestion reduced the yield from Saeman’s hydrolysis, but in this case did not increase the yields for the met/TFA hydrolysis. Our results do not support the use of Driselase digestion to increase saccharide yields.

**Table 4:** Saccharide composition of onion epidermal wall pretreated with chloroform and pre-digested with Driselase enzyme followed by met/TFA/Saeman’s hydrolysis.

| <u>µg/mg dry weight</u> |  |  |  |  |  |  |  |  |  |  |  |
| --- | --- | --- | --- | --- | --- | --- | --- | --- | --- | --- | --- |
|  | Trial 1 |  |  | Trial 2 |  | Trial 3 |  | mean |  | total | % |
|  | Chl+ Dri+<br>mTFA | Sae |  | Chl+ Dri+<br>mTFA | Sae | Chl+ Dri+<br>mTFA | Sae | Chl+ Dri+<br>mTFA | Sae |  |  |
| Glc | 41.0 | 63.4 |  | 20.9 | 9.0 | 97.6 | 6.9 | 53.2<br>(18.7) | 26.5<br>(15.1) | 79.6 | 19 |
| GalA | 190.3 | 22.7 |  | 208.3 | 4.9 | 287.4 | 26.9 | 228.7<br>(24.3) | 18.2<br>(5.5) | 246.8 | 60 |
| Gal | 32.3 | 4.2 |  | 14.7 | 0.1 | 84.6 | 2.2 | 43.9<br>(17.2) | 2.1<br>(1.0) | 46.0 | 11 |
| Xyl/Man* | 13.3 | 9.7 |  | 6.3 | 2.0 | 28.5 | 2.7 | 16.0<br>(5.4) | 4.8<br>(2.0) | 20.8 | 5 |
| Rha | 6.0 | 0.3 |  | 5.5 | 0.0 | 13.2 | 0.5 | 8.3<br>(2.0) | 0.3<br>(0.1) | 8.5 | 2 |
| Ara | 5.1 | 0.8 |  | 1.8 | 1.2 | 9.5 | 0.4 | 5.5<br>(1.8) | 0.8<br>(0.2) | 6.3 | 2 |
| Fuc | 1.4 | 1.7 |  | 0.0 | 0.1 | 3.8 | 0.0 | 1.7<br>(0.9) | 0.6<br>(0.5) | 2.3 | 0.6 |
| GlcA | 0.0 | 0.0 |  | 0.0 | 0.0 | 2.8 | 0.0 | 0.9<br>(0.8) | 0.0<br>(0.0) | 0.9 | 0.2 |
| **total** | **289.4** | **102.8** |  | **257.5** | **17.2** | **527.5** | **39.6** | **358.1** | **53.2** | **411.3** | **100** |

Chl+Dri+mTFA, peel is pretreated with chloroform and then Driselase digested. Soluble sugars are hydrolyzed by methanolysis with TFA
Sae, 2-step sulfuric acid (Saeman's hydrolysis) on the residue
Xyl/Man\*, HPAEC-PAD parameters did not separate xylose and mannose peaks
µg/mg values were adjusted using correction factors listed in Methods
(SEM), standard error of mean of 3 biological replicates

### GC/MS analyses

The foregoing results, based on met/TFA/HPAEC-PAD, were compared with GC/MS analysis of (a) per-*O*-trimethylsilyl (TMS) derivatives of the sugar methyl glycosides produced by acidic methanolysis (45) and (b) methyl alditols produced by permethylation, which required two different column separations (Figure 2) (46). The walls were prepared as above and extracted with chloroform to remove waxes, leaving cutin and unextractable waxes, so the yields are based on whole dewaxed walls.

**Figure 2:**
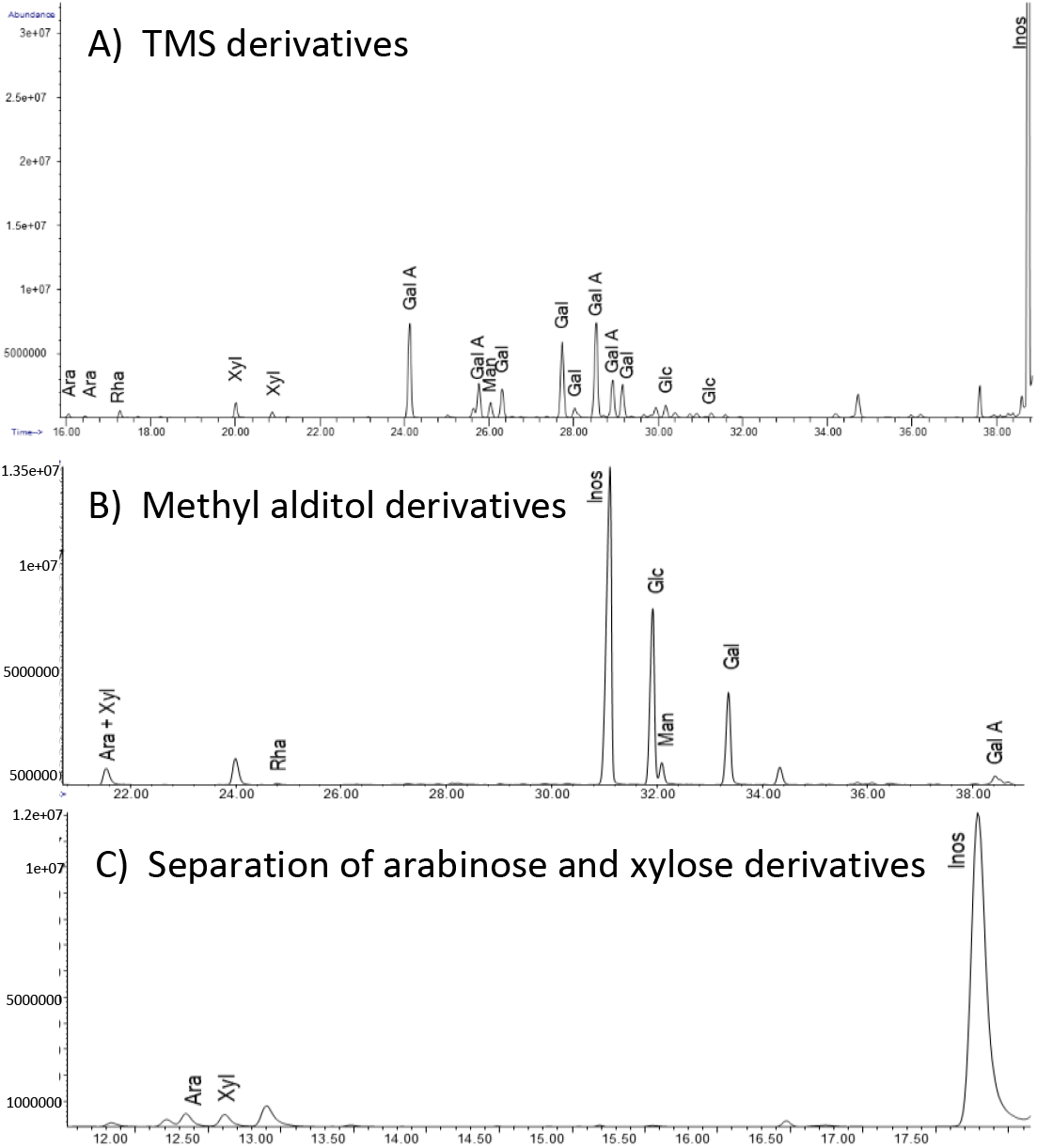
GC/MS chromatograms of chloroform pretreated onion epidermal peel using two different derivatization methods. **A**) TMS derivatives of the monosaccharide methyl glycosides separated on an Equity-1 fused silica capillary column; **B**) methyl alditol derivatives separated on an Equity-1 fused silica capillary column; **C**) same sample as **B** but run on a SP2330 column to separate arabinose and xylose derivatives.

The total yield by TMS methyl glycoside analysis (Table 5), was comparable to the best yield obtained with the met/TFA/S/HPAEC-PAD assays. GalA and galactose were the predominant sugars, but glucose and several other sugars were notably under-represented, compared with HPAEC-PAD results. The TMS method gives the highest yield for GalA (531 μg per mg of dewaxed wall) compared with the other results. In contrast, GC/MS analysis of methyl alditols showed substantial glucose content (223 μg per mg of dewaxed wall), but GalA was exceptionally low in the analysis. This method loses GalA but gives better yields for Glc and other neutral sugars. The GC/MS analyses find a 55:45 ratio of Xyl:Man, which we can use to estimate a Xyl value from Xyl/Man peak in the HPAEC-PAD analyses.

**Table 5:**
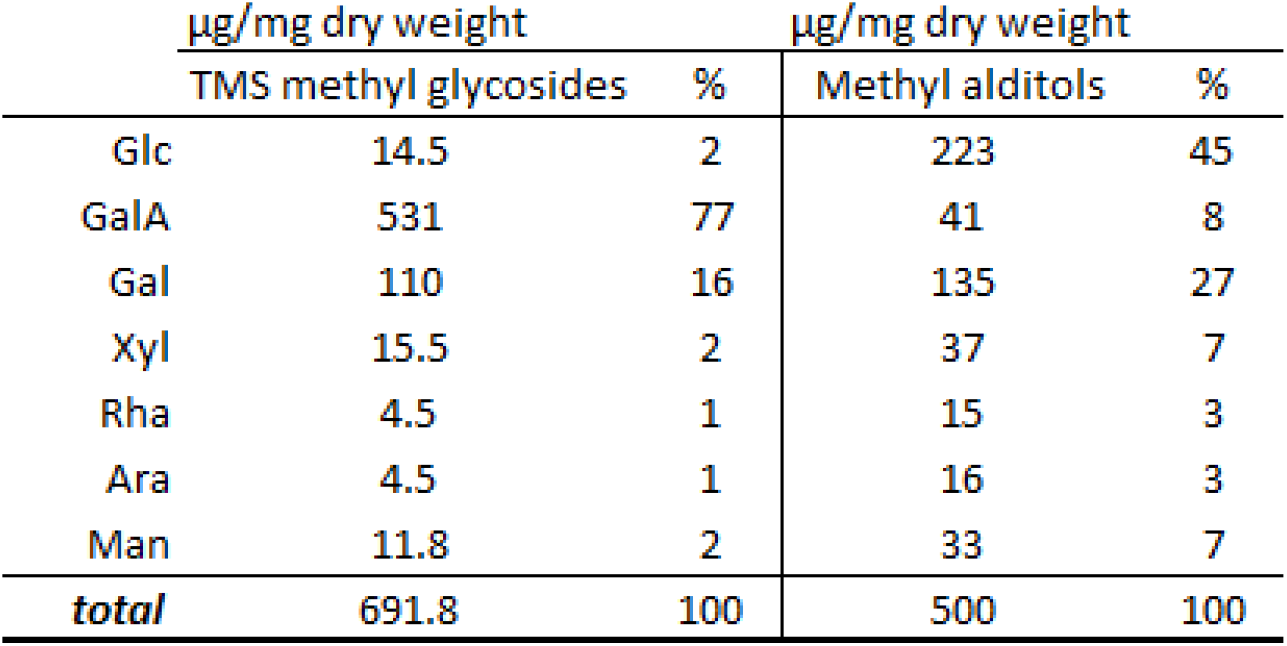
GC-MS analysis of onion outer epidermal walls derivatized to TMS methyl glycosides and methyl alditols.

| µg/mg dry weight |  |  | µg/mg dry weight |  |
| --- | --- | --- | --- | --- |
|  | TMS methyl glycosides | % | Methyl alditols | % |
| Glc | 14.5 | 2 | 223 | 45 |
| GalA | 531 | 77 | 41 | 8 |
| Gal | 110 | 16 | 135 | 27 |
| Xyl | 15.5 | 2 | 37 | 7 |
| Rha | 4.5 | 1 | 15 | 3 |
| Ara | 4.5 | 1 | 16 | 3 |
| Man | 11.8 | 2 | 33 | 7 |
| **total** | **691.8** | **100** | **500** | **100** |

Table 6 presents a comparative summary of the six preceding analyses, based on mol% of the total of measured monosaccharides in each dataset. The averaged HPAEC-PAD results show that Glc makes up a quarter of the monosaccharides, with twice as much GalA, half as much Gal and lesser amounts of other sugars. The GC/MS results were distorted by the severe loss of Glc in the TMS method and loss of GalA in the methyl alditol method. To compensate, we merged the two datasets by replacing the low GalA value of the latter method with the value from the TMS analysis, and recalculated mol% for this merged dataset (column G in Table 6). The composition of this merged data set is very similar to the averaged values in the HPAEC-PAD results.

**Table 6:** Comparative summary of monosaccharides in the onion epidermal wall (mol %)

| COLUMN: | HPAEC-PAD |  |  |  |  | GC/MS |  |  |
| --- | --- | --- | --- | --- | --- | --- | --- | --- |
|  | A | B | C | D | AVERAGE | E | F | G |
|  | mTFA/S | Chl + mTFA/S | Dri + mTFA/S | Chl/Dri+ mTFA/S |  | TMS | Methyl Alditols | Col E&F MERGED |
| Glc | 25 | 35 | 17 | 20 | 24 | 2 | 44 | 23 |
| GalA/GlcA | 49 | 43 | 58 | 58 | 52 | 75 | 7 | 51 |
| Gal | 14 | 11 | 14 | 12 | 12 | 17 | 27 | 14 |
| Xyl | 5 | 5 | 4 | 4 | 5 | 3 | 9 | 5 |
| Rha | 3 | 2 | 3 | 2 | 2 | 1 | 3 | 2 |
| Ara | 2 | 2 | 3 | 2 | 2 | 1 | 4 | 2 |
| Fuc | 1 | 1 | 1 | 1 | 1 | -- | -- |  |
| Man | 2 | 2 | 2 | 2 | 2 | 2 | 6 | 3 |
| **total** | **100** | **100** | **100** | **100** | **100** | **100** | **100** | **100** |

mol%, determined using 'total µg per mg' values for each experiment
Column G merges the GalA/GlcA µg per mg amount with the methyl alditol amounts
mTFA/S, methanolysis followed by TFA and Saeman's hydrolysis
Chl+mTFA/S, chloroform pretreated peel, methanolysis followed by TFA and Saeman's hydrolysis
Dri+mTFA/S, Driselase digested, methanolysis followed by TFA and Saeman's hydrolysis
Chl/Dri+mTFA/S, chloroform pretreat, Driselase digested, methanolysis followed by TFA and Saeman's hydrolysis

### Polysaccharide composition

We estimated the polysaccharide composition of the epidermal walls from the monosaccharide data, using an accounting method detailed in Materials and Methods. Homogalacturonan (HG) is by far the largest component at ~52% (by weight of total sugars); cellulose is estimated at 18-19% followed by galactan and xyloglucan at 11-13%. Other polysaccharides are present in amounts <5%. The last column in Table 7 includes relative polysaccharide content based on ^13^C solid-state NMR, to be discussed next.

**Table 7:**
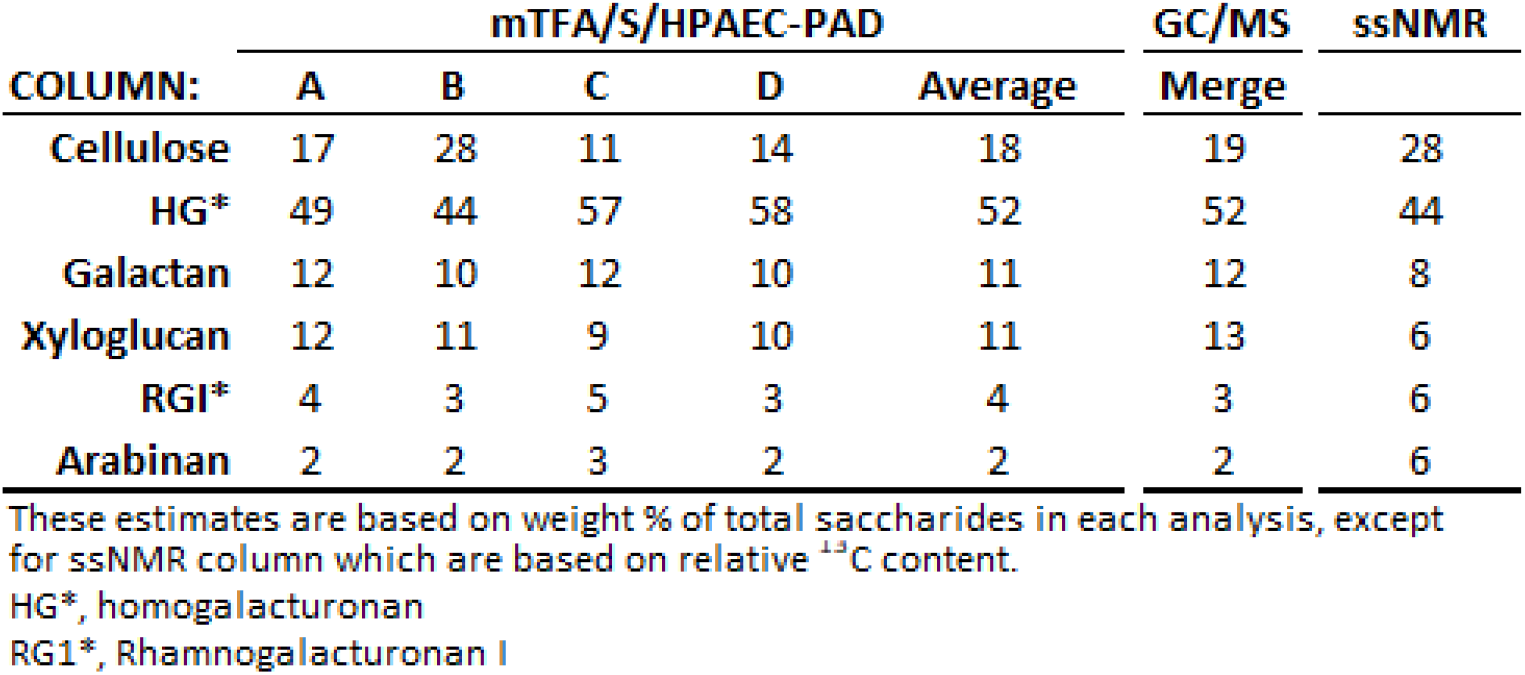
Comparative estimates of the relative polysaccharide composition of onion epidermal walls by different methods.

### Analysis by 1D ^13^C solid-state NMR (ssNMR)

A key advantage of this method is that cell walls do not need to be hydrolyzed, solubilized or otherwise chemically modified for ssNMR analysis, which measures all ^13^C atoms in the sample. The key disadvantage is that overlapping of ^13^C chemical shifts makes identification of many sugars difficult. This latter problem can be resolved in part by 2D and 3D spectral analysis, but the requirement of ^13^C enrichment was not feasible for our samples.

Epidermal walls were prepared as described above, extracted with chloroform to remove soluble waxes, rehydrated, and analyzed by two cross polarization (CP) methods. The conventional ^1^H-^13^C CP measurement, which selectively detects rigid components (Fig. 3a), showed major signals from the interior (i) and surface (s) glucan chains in cellulose microfibrils, such as the 89 ppm peak of interior cellulose carbon 4 (i4) and the 85 ppm peak of surface cellulose carbon 4 (s4). Additional major signals come from the GalA/GlcA carbon 1 (GA1) at 101 ppm, indicating pectin backbones are partially rigid in this sample. Using DMfit software (47), we performed spectral deconvolution of 1D ^13^C solid-state NMR spectra to quantify the polymer composition in the rigid phase of onion cell walls. The fit is of high quality, which is evidenced by the match in spectral patterns between the experimental and simulated spectra. The linewidths and areas (integrals) of all individual components are summarized in Additional file 1.

**Figure 3:**
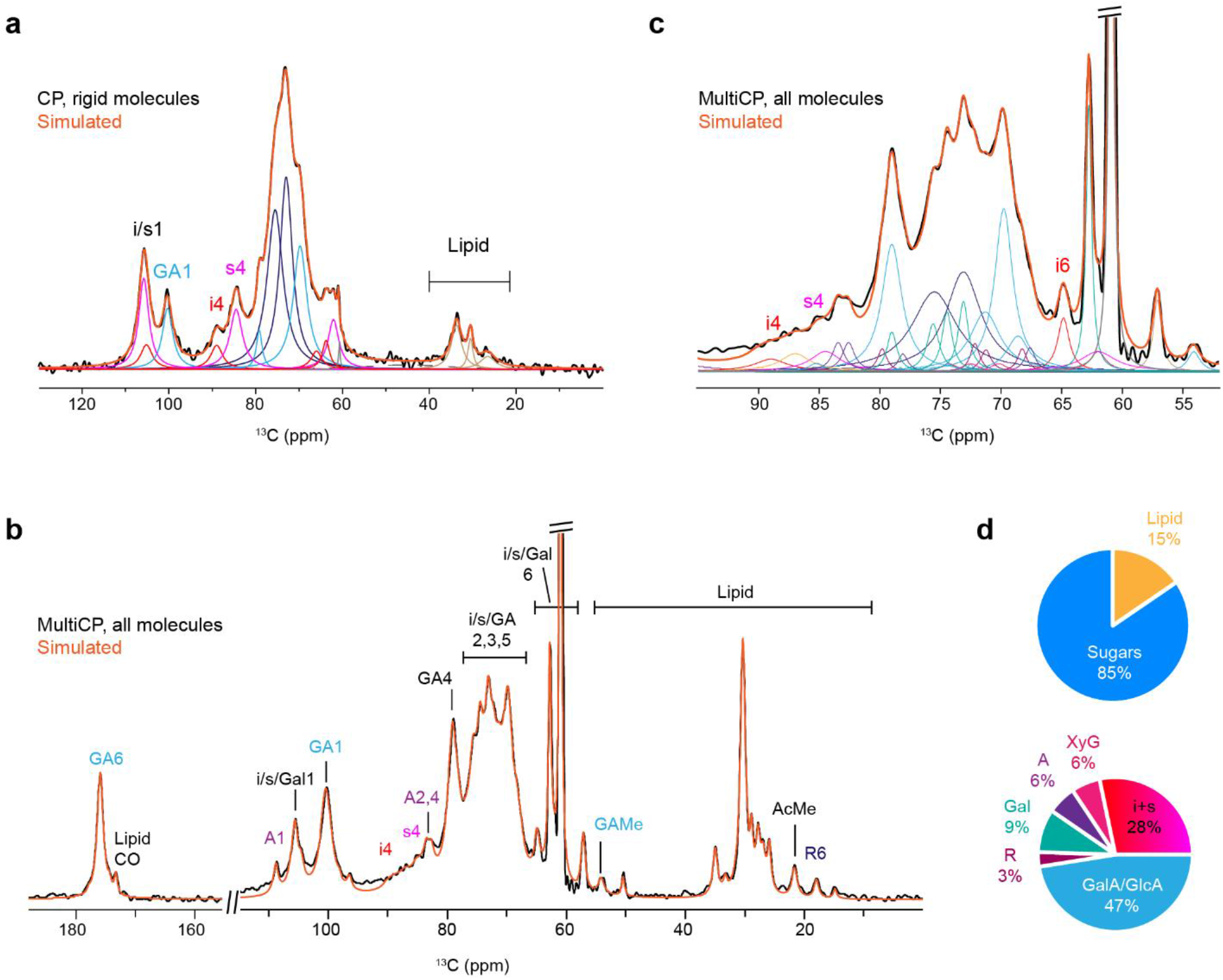
Solid-State NMR analysis of onion primary cell walls. **a,** Spectral deconvolution of 1D ^13^C conventional CP spectrum that mainly detects rigid components. The experimental data is in black and the fitted line is in orange. Underlying Lorentzians are individual fitted components. **b,** Spectral deconvolution of quantitative MultiCP spectrum that detects all molecules. **c,** Expanded region of 52-95 ppm region of MultiCP spectrum. **d,** Polymer composition from integrated ssNMR MultiCP spectra. Spectral deconvolution yields both polysaccharide to lipid ratio and sugar composition analysis. i (interior cellulose), s (surface cellulose), GA (GalA/GlcA), Gal (galactose), A (arabinose), R (rhamnose), GAMe (GalA/GlcA methylated), AcMe (acetylated methyl group), Lipid CO (carbon/oxygen), and XyG (xyloglucan). Following number refers to the associated ^13^C site number.

A second CP method made use of the recently developed MultiCP technique that detects all carbons in a quantitative manner (Fig. 3b,c) (48). The detailed procedures for achieving a satisfactory fit are described in Additional files 2 and 3. Briefly, cellulose peaks are first simulated using the information obtained from the CP spectrum, followed by (GalA), arabinose and rhamnose using their signature peaks, for example, carbon 1 of arabinose (A1) at 108 ppm and carbon 6 of rhamnose (R6) at 18 ppm. After all these carbohydrate components are fixed, two ambiguous components, the galactose and xylose are incorporated at the last stage. The reference chemical shifts of cell wall polysaccharides are obtained from the literature-reported high-resolution 2D ^13^C-^13^C correlation spectra collected using ^13^C-labelled plant materials (see (49) for a recent review), which are indexed in the Complex Carbohydrate Magnetic Resonance Database (CCMRD, (50)).

The polymer composition (by carbon number) is presented in Fig. 3d and Additional file 4. The sample contained 15% lipids and 85% carbohydrates. The lipid is likely to be from cutin, a cross-linked fatty acid polymer that is not removed by chloroform. Residual waxes may also contribute to the lipid peaks. Cellulose (i and s) accounts for approximately a quarter of all carbohydrates, while uronic acids are present in ~twice this amount. This conclusion is supported by the very low intensities of cellulose peaks (Fig. 3b), which is substantially lower than those observed in other plants such as *Arabidopsis* seedlings and maize primary cell walls (51, 52). Moreover, a major ambiguity exists in the quantification of xyloglucan backbone, whose signals severely overlap with the surface glucan chains of cellulose microfibrils and partially with the Gal signals in 1D spectrum as well. Consequently, the actual percentage of cellulose and galactose might be slightly lower than the data presented in Fig. 3d (see Additional file 5).

In comparing the ssNMR results with those of the ‘wet-bench’ methods (summarized in Table 7), we find generally good agreement except that cellulose is higher and xyloglucan lower in the ssNMR results. This discrepancy could be partly due to difficulty in differentiating xyloglucan signals from cellulose in the ssNMR spectra and partly the result of losses of cellulose in the wet-bench methods.

## Discussion and Summary

From these collective results we estimate the onion outer epidermal wall to be composed predominantly of pectic polysaccharides (~50% homogalacturonan, ~10% galactan, ~5% rhamnogalacturonan I, and 2-6% arabinan, as weight % of total sugars), with ~20% cellulose and smaller amounts as xyloglucan (~10%). Variations in quantitation and yields seen with the different methods limit the precision of these values and highlight the importance of reporting yields as a percentage of the dry wall mass as well as percentage of total sugars. Total yield is often omitted from published studies of cell wall compositions. Yields, evaluated as μg sugar per mg dry weight of wall and obtained with ‘wet-bench’ methods, were less than the maximum sugar content (85%) determined by ^13^C ssNMR of the de-waxed epidermal wall, which contained 15% residual lipid. This comparison is biased by two factors that influence the wet-bench yields: (a) The lipid content of the epidermal wall reduces the calculated yields. (b) The addition of water to the glycosyl residues upon hydrolysis exaggerates the yields based on summing the monosaccharides, by ~10%. To illustrate the magnitude of these two combined effects, if we adjust the average yield reported in Table 2 for these two factors, the yield increases from 725 to 768 μg per mg. This approaches the ssNMR estimate of 85% sugar based on MultiCP spectra. If we include the waxes removed by chloroform (~12% of the wall dry weight), we estimated that ~one-quarter of the onion outer epidermal wall mass is lipid. The ssNMR estimate of 85% of saccharide content is based on ^13^C only and takes into consideration the non-extractable lipids in the wall but does not include inorganic substances in the wall, which have not been evaluated. Metal ions bound to the pectins and silica will reduce gravimetric-based assessments of sugar yields. Finally, sugar losses that are inherent in analytical methods reliant on chemical hydrolysis and derivatization (35, 40, 46) may be aggravated in the epidermal wall by its impregnation with hydrophobic substances (53, 54) which may hinder polysaccharide solubilization and hydrolysis.

These results with the onion scale epidermal wall are roughly comparable to sugar analyses of cell walls from onion parenchyma which likewise document a pectin-rich cell wall dominated by homogalacturonan and galactan with smaller amounts of xyloglucan and cellulose (21–24). However, differences in the onion materials and analytical methods make quantitative comparisons difficult. There are few published analyses of the saccharide composition of epidermal walls. One such study, of growing maize coleoptiles (55), reported the epidermal wall to contain more cellulose (59% of wall sugars) and less matrix polysaccharides (39%) compared with mesophyll walls (28% and 72%, respectively). Lipid content of the epidermal wall was not measured in this study. Nevertheless, it seems clear that outer epidermal walls of onion scales contain much less cellulose than those of the maize coleoptile, if these two studies are representative of the two materials; additionally, the striking difference between epidermis versus mesophyll parenchyma reported for maize coleoptiles does not appear to be the case for onion scales. Saccharide analyses of other epidermal walls would be useful to characterize the natural diversity in this feature and its relationship to epidermal functions.

To confirm our ‘wet bench’ (HPAEC/PAD and GC/MS) results, we used 1D NMR to examine the walls without the need for solubilization and depolymerization, which entail variable losses because of incomplete hydrolysis and chemical degradation of sugars. It remains technically difficult to resolve glucose in xyloglucan and cellulose by ssNMR; therefore, coupling solid-state NMR with other biochemical and bioanalytical assays is a promising strategy for investigating plant cell walls. The CP spectrum provides reference peaks for initiating the simulation of the more complicated MultiCP spectrum that contains a larger number of components (Fig. 3b, Additional file S3). This method allows us to overcome the challenges associated with the limited resolution of a 1D NMR spectrum due to overlapping peaks (Fig. 3c). Moreover, it provides consistency of peak width and amplitude (and so integral) within each individual cell wall constituent.

We achieved a good match between the experimental and simulated spectra, and the general coherence in surface of individual components within a single carbohydrate component. In addition, the partially resolved lipid peaks are also included in the simulation. The recent development of natural-abundance Dynamic Nuclear Polarization and its application to biomass materials allow us to collect high-resolution 2D correlation spectra using unlabeled materials at natural isotope abundance (1% for ^13^C), which will further improve the accuracy of ssNMR analysis (56–58).

## Conclusion

The outer epidermal wall of onion scales was estimated to contain ~50% homogalacturonan (as % of total wall saccharide), ~20% cellulose, ~10% galactan, ~10% xyloglucan and smaller amounts of other polysaccharides. The estimate is based on chemical and enzymatic hydrolysis products by HPAEC-PAD, GC/MS analysis of methyl alditols and per-*O*-trimethylsilyl derivatives, and 1D NMR of whole walls. Our results illustrate method-dependent variations in total yields and indicate that up to one quarter of the dry matter of the epidermal wall may be non-extractable cutin and extractable waxes that may interfere with saccharification.

## Materials and Methods

### Fresh onion preparation

Commercial white onions, 8 – 10 cm in diameter, were purchased from a local grocery store. The onion dry skin was removed and a rectangular cut was made to remove the sections of onion. The 5th layer from first outside fleshy layer was used for all experiments. For each experimental replicate, one whole onion’s 5th layer was harvested. The 5th layer was peeled by hand and forceps by snapping off one of the ends of the scale and gently peeling back the outer (abaxial side) epidermal layer of cells. This layer will be very translucent and thin, as it only contains half of the epidermal wall, cutin layer, and outer wax layer. The peels were laid flat with cuticle side up, the extra flesh ends were cut off with a razor blade, and the peels were floated in a petri dish of water to briefly wash. The peels were submerged in 2% solution (w/v) of sodium dodecyl sulfate (SDS; Sigma, USA) in a 50 mL Falcon tube. Tubes were rotated end-for-end for 4 h. The SDS solution was saved, water added to the tubes, and vortexed briefly to wash out SDS. A final wash was done by retaining the peels over a 40 μm cell strainer and then pouring more water over the peels until no further bubbles could be seen.

The SDS washed peels were then de-starched by incubation with α-amylase from porcine pancreas (Sigma, USA) without losing additional wall polymers (59). 100 U of amylase powder was added to 50 mL of 25 mM TRIS-HCL, pH 7.5 with 100 mM KCl added, incubated with shaking for 30 min, then filtered through a 0.45 μm PES syringe filter. The filtered solution was added to onion peels, 2 mM sodium azide was added to control bacterial growth, and tube was rotated end-over-end for 16 h at room temperature. Amylase enzyme was removed by washing the peels with 2% SDS (3x exchange), followed by ddH_2_O wash (3x exchange) to remove the SDS. Peels were then put in a 50 mL Falcon tube with enough water added to cover them, the cap had holes poked in it for vapor exchange, and the tubes frozen in – 80°C for at least 2 h, then lyophilized overnight. Dry peels were then weighed (weight post SDS and amylase). These peels were used for all subsequent characterizations.

### Methanolysis / TFA hydrolysis

SDS washed and de-starched dry onion peels were weighed and added to a 2 mL screw – top microcentrifuge tube and 1 mL of 3 M HCl in methanol (Sigma, USA) was added to the walls and incubate for 16 h at 80°C in heating block in a fume hood as per De Ruiter et al, 1992. The tubes were cooled on ice, briefly spin down to collect condensate, and evaporated under a gentle stream of airflow at 25°C. One mL of 2 M TFA stock (Sigma, USA) was added to the methanolic HCl-treated walls and incubated for 1 h at 120°C, cooled down on ice, and centrifuged at 8,000 g for 5 min to pellet any solids. The supernatant from the met/TFA hydrolysis was separated from the remaining residue using a pipette and both fractions were saved in separate tubes.

Both tubes had any remaining liquid evaporated under a gentle stream of airflow at 25°C. The dried soluble supernatant was then resuspended in 1 mL ddH20 and filtered through a 0.2 μm nylon filter for anion exchange chromatography. The insoluble residue had a small amount of water added to the tube, frozen in – 80°C, and lyophilized overnight. To correct for monosaccharide losses due to degradation during the met/TFA hydrolysis, the μg per mg amounts for each monosaccharide (Tables 1 – 4) had correction factors applied according to (35): Glc (0.94), GalA (0.71), Gal (0.86), Xyl (0.91), Rha (0.83), Ara (0.83), Fuc (0.93), GlcA (0.63), Man (0.89).

### Saeman’s hydrolysis

The lyophilized residue after met/TFA hydrolysis was treated with a 2-step sulfuric acid protocol to break down crystalline cellulose remaining with minor changes to the Saeman’s protocol (43). Briefly, the dried residue was added to a new 2 mL screw- top microcentrifuge tube and 100 μl of 72% (w/w) H_2_S0_4_ was added. The tube was shaken at 25°C until the residue dissolved or became translucent (usually less than 3 h; in some cases the residue never completely dissolved). Nine hundred μl of ddH20 was added to the solution and mixed well with inversion. The tube was heated at 120°C for 1 h. The tube was cooled on ice and then neutralized using 6M NaOH. The solution was then filtered through a 0.2 μm PVDF syringe filter to remove any residue remaining (if large insoluble material remained, the insoluble residues was sedimented at 10,000 g for 10 min and then filtered to collect the supernatant). Samples were stored in −20 °C until time for analysis. To correct for monosaccharide losses due to degradation during the Saeman’s hydrolysis, the μg per mg amounts for each monosaccharide (Tables 1 – 4) had correction factors applied according to (60) for Gal, Xyl, Ara, Rha, and Man, (21, 60) for the recovery of glucose, and (61), for the recovery of GalA: Glc (0.49), GalA (0.5), Gal (0.94), Xyl (0.82), Ara (0.96), Rha (0.93), Man (0.87).

### Chloroform treatment

SDS washed and de-starched dried onion peels (20 mg) were rehydrated in 5 mL chloroform in a glass screw top tube for 1 h with solution stirring using a micro stir bar, then fresh chloroform was exchanged for 24 h incubation with stirring at room temperature. The chloroform supernatant was removed using a pipette and put into a separate glass tube and evaporated with air; the peels were also dried down with air, frozen in ddH_2_O, lyophilized, and re-weighed. The peels were then processed by met/TFA hydrolysis to obtain the soluble sugar fraction and the residue from this hydrolysis was sequentially treated by Saeman’s hydrolysis.

### Driselase digestion

SDS-washed and de-starched dry onion peels were weighed and put in a 5 mL screw-top conical tube containing 5 mL of 20 mM sodium acetate buffer, pH 5.5 with 2 mM sodium azide. Twenty milligrams (20 mg) of peels were used and 50 μL of a stock of Driselase enzyme (nominally 10 mg/mL) was added to the tube. The Driselase stock solution was made by dissolving 10 mg of Driselase powder (Sigma, USA) in 1 mL of ddH_2_O and shaking for 30 minutes at 800 rpm. The tube was then spun down at 10,000 rpm for 5 minutes and the supernatant was collected and filtered through 0.2 μm PVDF filter. Sodium azide was added to the tube (2 mM final concentration) to reduce microbial growth. Peels were incubated at 37°C and rotated end-for-end for 7 days. Tubes were then centrifuged at 10,000 rpm for 10 minutes and the supernatant was collected by pipette and filtered through a 0.2 μm nylon filter. The peel residue was washed with ddH_2_O two times to remove the sodium acetate buffer. Both soluble and insoluble fractions where then frozen in water and lyophilized overnight. Both the enzyme supernatant and the peel residue were then processed by met/TFA hydrolysis and Saeman’s hydrolysis, respectively.

### High-Performance Anion-Exchange Chromatography with Pulsed Amperometric Detection

All dried supernatants were resuspended in 1 mL of ddH_2_O, incubated at 25°C for 30 minutes, and filtered through a 0.2 μm nylon filter (VWR, USA). Enzyme-treated samples were first centrifuged through a 3-kDa spin column (Nanosep, USA) to remove any trace of enzyme while monosaccharides are eluted. The samples were then diluted 1:10 in ddH_2_O for the enzyme and met/TFA hydrolysates, and 1:20 for the Saeman’s hydrolysates, respectively. We used a Dionex ICS5000 single pump chromatography system (Thermo Fisher Scientific, USA) equipped with pulsed amperometric detection with disposable gold electrodes, and a WSP 3000 autosampler (Thermo Fisher Scientific, USA). The CarboPac PA20 column (Thermo Fisher Scientific) was kept at 30°C during the injections and all eluents were filtered through 0.2 μm PES filter (VWR, USA) and stored under helium. The chromatography conditions were as follows: equilibration in 90% ddH_2_O + 10% of 100 mM sodium hydroxide for 5 minutes, then in 90% ddH_2_O + 10% of 100 mM NaOH for 15 minutes after injection of sample to elute neutral monosaccharides, followed by a 0 – 100% gradient of [100 mM sodium acetate + 100 mM sodium hydroxide] in 100 mM sodium hydroxide for 25 minutes to elute acidic monosaccharides, and a final column wash of 200 mM sodium hydroxide for 5 minutes. The flow rate was 0.5 mL/min and injection volumes were 10 μL for all runs. Monosaccharide standards, unhydrolyzed, were commercially sourced (Sigma, USA) and injected at 10 μg/mL concentration in ddH_2_O in triplicate. Three injection replicates were averaged for compositional analysis, and all experiments had three biological replicates which are reported in the tables. Chromeleon software v 7.3 (Thermo Fisher Scientific) was used to create reports and export the raw data to Excel for further statistical analysis.

### Glycosyl composition of TMS derivatives

Glycosyl composition analysis was done using gas chromatography/mass spectrometry (GC/MS) of the per-*O*-trimethylsilyl (TMS) derivatives of the monosaccharide methyl glycosides produced from the sample by acidic methanolysis (45). The chloroform-treated whole peel (110 μg) was heated with methanolic HCl for 17 h at 80°C in a screw-top glass test tube. The sample had particulates floating in the solution after methylation, potentially cuticular material. The samples were cooled and the liquid was dried off using a stream of nitrogen. The tube was then treated by adding a mixture of methanol, pyridine, and acetic acid anhydride for 30 minutes. Solvent was evaporated and the sample was derivatized using Tri-Sil (Pierce, USA) at 80°C for 30 minutes. TMS methyl glycosides were then analyzed on an Agilent 7890A GC interfaced to a 5975C MSD using an Equity-1 fused silica capillary column (30 m × 0.25 mm ID, Supelco, USA).

### Glycosyl composition of methyl alditol derivatives

Chloroform-treated whole peels (100 μg) were permethylated by two rounds of treatment with sodium hydroxide (15 minutes each) and methyl iodide (45 minutes each) in DMSO (Sigma, USA) (46). The onion peel sample was then hydrolyzed by adding 2 M TFA to the tube and heated for 2 h at 121°C, reduced with NaBD4, and remethylated; the second methylation involved a single treatment with base and methyl iodide. The resulting methylated alditols were analyzed on the same instrumentation and column as the TMS methyl glycosides. Separation of the xylose and arabinose residues required an additional run on a SP-2331 bonded-phase fused-silica capillary column (Supelco, USA).

### Solid-state Nuclear Magnetic Resonance

The unlabeled chloroform-treated onion cell walls (107 mg wet mass) were packed into a 4-mm magic-angle spinning (MAS) rotor for measurements on a 400 MHz Bruker Avance solid-state NMR spectrometer. Two types of experiments were conducted: the conventional ^1^H-^13^C cross polarization (CP) that selectively detects rigid molecules and the recently developed MultiCP technique that enables efficient detection of all molecules using unlabeled materials in a quantitative manner (48). Both experiments were conducted at room temperature under 14.0 kHz MAS. In total, 24,576 scans were recorded for the CP spectrum and 42,944 scans were collected for the MultiCP spectrum. The detailed procedures for spectra deconvolution are provided in the Additional Flies.

### Estimation of polysaccharide ratios from monosaccharide composition

Our procedure for estimating polysaccharide contents from sugar analysis included the following sequential steps:

1. Xyloglucan: We assign all Xyl to xyloglucan and then assign corresponding amounts of Glc, Gal and Fuc to xyloglucan based on the ratio Glc:Xyl:Gal:Fuc (5:3:1:0.5) from an analysis of onion xyloglucan (26).
2. Cellulose: The remaining Glc is assigned to cellulose.
3. Galactan: All the Gal not assigned to xyloglucan is assigned to galactan.
4. RGI: All the Rha is assigned to RGI along with equimolar amounts of GalA, corrected for the molecular weight difference of GalA.
5. HG: All the remaining GalA remaining after the RGI assignment is assigned to HG.

## List of abbreviations

ddH_2_O: double distilled water
HPAEC-PAD: high-performance anion-exchange chromatography with pulsed amperometric detection
H_2_SO_4_: sulfuric acid
NaOH: sodium hydroxide
NaOAc: sodium acetate
PES: polyethersulfone
PVDF: polyvinylidene difluoride
SDS: sodium dodecyl sulfate
TFA: trifluoracetic acid
TMS: trimethylsilyl
ssNMR: solid-state nuclear magnetic resonance

## Declarations

### Authors’ contributions

LW performed all hydrolysis and anion exchange chromatography experiments, collected data, analyzed data, made figures, and made a rough draft of the manuscript. DC oversaw all experimental design and results, editing of the manuscript, and sharing of results within our group. TW/FD performed all ssNMR experiments, collected data, analyzed ssNMR data, and prepared ssNMR figures. All authors read and approved final manuscript.

## Acknowledgements

We thank Drs. Ian Black and Parastoo Azadi at the Complex Carbohydrate Research Center for the GC-MS analysis of the epidermal samples and discussion of the results. We also thank Dr. Sarah Kiemle for initial trials with the methanolysis with TFA protocol, and Edward Wagner for the Driselase protocol and protein determination.

## Ethics approval and consent to participate

Not applicable

## Consent for publication

Not applicable

## Availability of data and materials

All data generated during this research study are included in this published article and its supplementary information files.

## Additional files

**Additional file 1. Table S1.** Parameters used for the fit of ^13^C CP spectrum in Figure 3a.

**Additional file 2.** Procedures for spectral deconvolution

**Additional file 3. Table S2.** Parameters used for the fit of ^13^C MultiCP spectrum in Figure 3b, c. The relatively well resolved resonances (underlined) are used for estimating polysaccharide composition.

**Additional file 4. Table S3.** Molecular composition derived from ^13^C MultiCP spectrum in Figure 3b, c. The relatively well resolved resonances (underlined) are used for estimating polysaccharide composition. Carbohydrate peaks account for 85% of total intensity and lipid polymers account for 15% of all carbons.

**Additional file 5. Figure S1.** MultiCP integration models.

