## Additional file for "Saccharide Analysis of Onion Outer Epidermal Walls"

**Additional file 1. Table S1.** Parameters used for the fit of ^13^C CP spectrum in Figure 3a.

| **δ [ppm]** | **Assignment** | **Amplitude** | **Width [ppm]** | **Integral [%]** |
| --- | --- | --- | --- | --- |
| Cellulose | | | | |
| 105.8 | i1 | 12 771 | 2.8 | 7.8 |
| 105.2 | s1 | 3 266 | 3.0 | 2.2 |
| 89.0 | i4 | 3 329 | 3.3 | 2.4* |
| 84.5 | s4 | 8 336 | 3.5 | 6.3* |
| 75.5 | i3 and s3/5 | 22 105 | 4.4 | 21.1 |
| 73.0 | i2/5 and s2 | 26 779 | 3.5 | 20.4 |
| 66.0 | i6 | 2 500 | 3.0 | 1.6 |
| 62.0 | s6 | 6 900 | 3.0 | 4.5 |
| Other major peaks | | | | |
| 100.3 | GalA1 | 8 506 | 2.9 | 5.4 |
| 79.2 | GalA4 | 5 449 | 1.6 | 1.9 |
| 69.8 | GalA3 | 17 283 | 3.8 | 14.2 |
| 63.8 |  | 4 000 | 2 | 1.7 |
| 61.0 |  | 3 900 | 0.4 | 0.3 |
| Other minor peaks | | | | |
| 45.4 |  | 631 | 25.6 | 3.4 |
| 33.7 |  | 6 165 | 2.9 | 3.8 |
| 30.5 |  | 4 336 | 1.4 | 1.3 |
| 26.5 |  | 1 821 | 4.3 | 1.7 |

* highlights values used to calculate the interior-to-surface ratios for cellulose, which is later used in Method 2 for quantifying polysaccharide composition in **Additional files 3** and **4**.

**Additional file 2. Procedures for spectral deconvolution**

Here is described the methodology employed to obtain complex ^13^C NMR spectra deconvolution presented in **Figure 3**. It should be noted that the peak position, shape, width, amplitude can all affect the result; therefore, prior knowledge of the representative linewidth and chemical shifts are a necessity to achieve a satisfactory fit. The representative chemical shift information of cell wall polysaccharides can be accessed at the Complex Carbohydrate Magnetic Resonance Database (CCMRD) at [www.ccmrd.org](http://www.ccmrd.org) [1].

**Step-by-step building of the simulated spectra**

First, deconvolution of the basic Cross Polarization (CP) spectrum (**Figure 3a**) is performed to obtain references and information on the chemical shift, linewidth, and intensity of cellulose peaks. It is straightforward to get a reasonable fit of the CP spectrum because only a small number of components are present. The information is then used to guide the fit of cellulose peaks in the quantitative MultiCP spectrum. In Dmfit software, the CP fit is directly loaded in the MultiCP experimental spectrum. After baseline correction, a normalization factor of 1.1 is applied to cellulose to match intensity levels of i4 and s4 peaks. Resulting cellulose amplitudes are locked, alongside chemical shifts and widths.

For CP fit convergence, 5 other major peaks and 4 minor ones are mandatory. 3 out of the 5 supplementary major peaks correspond to the most abundant constituent of the onion cell wall, galacturonic acid. We note that this assignment of CP spectral lines to GalA and cellulose is in good agreement with past literature on major polysaccharides found in onion cell wall. Then, complete MultiCP fit (**Figure 3c**) is built up by adding components, one sugar after the other. Final parameters are given in **Additional file 3**.It began with arabinose and rhamnose, as they both display a well resolved carbon (respectively A1 and R6) that can be fitted without overlap of other unknown sugars at this this stage. Resulting amplitudes and widths from these A1 and R6 sites are applied to all other arabinose and rhamnose carbons. It must be noted that for A1, a visual fit was done (in other words, parameters for A1 were manually adjusted; the ‘compute’ command in DmFit was not used). It is followed with galactose (G6), for which data is obtained after local fit of the 0 to 50 ppm region, which includes its resolved peak around 18 ppm. Then, to fill up the 68 to 80 ppm region (chemical shift interval where spectral overlap is too great to distinguish any resolved peak), results of the 160 to 180 ppm local fit are used. The latter is considered a good starting point, as it requires only 3 lines, including 2 attributed to the components of interest, GalA and GlcA. Once again, amplitude and widths are applied to all other carbon of GalA and GlcA. At this stage, visual inspection clearly shows that overall line shape requires corrections, as several amplitudes of these sites are overestimated. Thankfully, it is noticed that major improvement of the fitted line shape can be obtained with adjustment of some originally CP-given spectral lines. (This may be justified, as local fit reference of C1 of other main constituent GalA/GlcA give anyway different widths of C3 and C4 compared to CP spectrum fit.) Furthermore, i1 cellulose peak width adjustment is operated, allowing the introduction of the last major sugar, galactose, through its C1. This way, minor contribution of xylose (at 104.5 ppm) is also taken in account (no other chemical shift area could have given fit parameters for this sugar).

Automatic fit computation, with only amplitude and widths of major components (about 20 parameters) allowed to vary, does not converge to a satisfactory result, even after an hour of calculations. Therefore, a manual fit is required. Parameters adjustments are generally made from low to high field to minimize extent of changes. After obtaining a satisfactory fit, fine tuning of C4 region is performed to discriminate the different cellulose polymorphs. With this model they are assumed to yield similar broad components as in CP spectrum. Without ^13^C labelling, considered cellulose conformers are interior chains, hydrophilic and hydrophobic surface chains, respectively referred by i, s^f^ and s^g^ in the literature [2].

**Estimate molecular fraction using deconvolution data**

From all deconvoluted spectral lines (including those classified as others in **Additional file 3**), we add all integrals for polysaccharides and lipid. Ratio of one to another yield 85% sugars to 15% lipid (**Additional file 4**). To obtain polysaccharide ratios from one to another, we mostly average selected integrals from peaks that are clearly resolved. Practically, this generally correspond to consider numerical integrals from the same peaks from which the fit has been initiated. Then, these averages are divided to the total amount of polysaccharides. Obtained ratios are used to establish charts presented in **Figure 3**. Formulas to obtain them are explicated below Table S2. Exception is made for cellulose. Indeed, to alleviate uncertainties, we propose 2 integration models. In the first one (presented in **Figure 3 and Figure S1**), all integrals from broad C4 peaks of cellulose are added, resulting in considering i4 and s4 as a representative amount of the total cellulose content in onion cell wall. This method yields a trustworthy cellulose content but does not allow us to discriminate interior from surface cellulose. In a second model, in order to try to get conformer specific information, we use resolved i6 peak as the reference and infer total cellulose amount by adding the hypothetical s6 contribution equaling i6 integral modulated by i to s ratio inferred from classic CP experiment.

Estimating the error is quite challenging. We expect around 10% error margin of each reported value based on the current signal-to-noise ratios. Additional error margin should be added to account the uncertainty due to the manual fitting of the spectrum. Therefore, we are confident on discriminating major components (GalA, cellulose) from minor ones (arabinose, rhamnose, xyloglucan), but acute proportions within each category could be subject to significant variation, especially coming from the broader areas of the spectrum.

**Additional file 3. Table S2.** Parameters used for the fit of ^13^C MultiCP spectrum in Figure 3b, c. The relatively well resolved resonances (underlined) are used for estimating polysaccharide composition.

| **δ [ppm]** | **Carbon number** | **Amplitude** | **Width [ppm]** | **Integral [%]** | **Contribution**  **Before normalization** | **Normalized fraction,**  **Method 1** | **Normalized fraction,**  **Method 2** |
| --- | --- | --- | --- | --- | --- | --- | --- |
| **GalA/GlcA** | | | | | | | |
| Molecule 1 | | | | | | | |
| 176.0 | 6 | 38 944 | 1.2 | 4.5 | 6.3 **^a^** |  |  |
| 100.3 | 1 | 32 000 | 2.2 | 6.6 |  |  |  |
| 79.0 | 4 | 32 000 | 2.0 | 6.0 |  |  |  |
| 71.3 | 5 | 15 000 | 3.0 | 4.2 |  | 47% | 42% |
| 69.8 | 3 | 41 000 | 1.8 | 6.9 |  |  |  |
| 68.6 | 2 | 9 000 | 2.2 | 1.9 |  |  |  |
| 54.1 | Me | 5 000 | 1.2 | 0.6 |  |  |  |
| Molecule 2 | | | | | | | |
| 177.9 | 6 | 943 | 0.6 | 0.1 | 0.1 |  |  |
| 98.1 | 1 | 1 800 | 0.8 | 0.1 |  |  |  |
| 78.3 | 3 | 943 | 0.5 | 0.1 |  | 0.7% | 0.7% |
| 72.5 | 2/5 | 1 886 | 0.6 | 0.1 |  |  |  |
| . | 4 | . | . | . |  |  |  |
| **Rhamnose (Rha)** | | | | | | | |
| 96.4 | 1 | 4 000 | 0.9 | 0.4 | 0.4 **^b^** | 3.0% | 2.8% |
| 79.8 | 2 | 5 730 | 0.9 | 0.5 |  |  |  |
| 72.2 | 4 | 7 000 | 0.9 | 0.6 |  |  |  |
| 71.3 | 3 | 5 450 | 1.1 | 0.5 |  |  |  |
| 68.3 | 5 | 5 730 | 0.9 | 0.5 |  |  |  |
| 17.8 | 6 | 5 730 | 0.9 | 0.5 |  |  |  |
| **Cellulose (i/s)** | | | | | | | |
| 105.7 | i1 | 11 500 | 1.7 | 1.8 | Method 1:  3.7 **^c^**  Method 2:  5.2 **^d^** | 28% | 35% |
| 90.0 | i^e^4 | 1 000 | 2.0 | 0.2 **^c^** |  |  |  |
| 89.0 | i^a/b^4 | 3 000 | 3.3 | 0.9 **^c^** |  |  |  |
| 87.0 | i^c/d^4 | 4 400 | 3.0 | 1.2 **^c^** |  |  |  |
| 84.5 | s4 | 5 000 | 3 | 1.4 **^c^** |  |  |  |
| 75.5 | i3 + s3/5 | 20 000 | 4.4 | 8.2 |  |  |  |
| 73.1 | i2/5 + s2 | 25 000 | 3.5 | 8.2 |  |  |  |
| 64.9 | i6 | 13 500 | 1.2 | 1.5 **^d^** |  |  |  |
| 60.9 | s6 | 5 000 | 3.5 | 1.6 |  |  |  |
| **Galactose (Gal)** | | | | | | | |
| 105.5 | 1 | 9 900 | 1.0 | 0.9 | 1.3 **^e^** | 9.3% | 8.3% |
| 79.0 | 4 | 9 900 | 1.0 | 0.9 |  |  |  |
| 75.6 | 5 | 12 000 | 1.2 | 1.4 |  |  |  |
| 74.5 | 3 | 15 000 | 1.0 | 1.4 |  |  |  |
| 73.1 | 2 | 17 500 | 1.0 | 1.6 |  |  |  |
| 62.8 | 6 | 67 000 | 0.6 | 3.5 |  |  |  |
| **Arabinose (Ara)** | | | | | | | |
| 108.7 | 1 | 9 200 | 1.1 | 1.0 | 0.9 **^f^** |  |  |
| 83.5 | 4 | 7 300 | 1.2 | 0.8 |  |  |  |
| 82.6 | 2 | 7 300 | 1.2 | 0.8 |  | 6.4% | 5.8% |
| 78.1 | 3 | 4 500 | 1.0 | 0.4 |  |  |  |
| 67.6 | 5 | 6 000 | 1.0 | 0.6 |  |  |  |
| **Xylose in xyloglucan (Xyl)** | | | | | | | |
| 99.6 | 1 | 2 000 | 1.6 | 0.3 | 0.3 |  |  |
| 74.3 | 3 | 2 000 | 1.6 | 0.3 |  |  |  |
| 72.5 | 2 | 2 000 | 1.6 | 0.3 |  | 2.2% | 2.0% |
| 70.2 | 4 | 2 000 | 1.6 | 0.3 |  |  |  |
| 62.0 | 5 | 5 000 | 1.6 | 0.8 |  |  |  |
| **Glucose in xyloglucan (Glc)** | | | | | | | |
| 104.5 | 1 | 7 100 | 1.2 | 0.8 | 0.5 ^g^ | 4.0% | 3.6% |
| 85.2 | 4 | 5 000 | 1.6 | 0.3 |  |  |  |
| **Other signals of Polysaccharides** | | | | | | | |
| 174.4 |  | 2 817 | 0.8 | 0.2 |  |  |  |
| 60.9 |  | 220 000 | 0.4 | 8.2 |  |  |  |
| 21.6 | Ac^Me^ | 9 137 | 1.1 | 0.9 |  |  |  |
| **Lipid** | | | | | | | |
| 173.3 |  | 5 925 | 0.6 | 0.3 |  |  |  |
| 57.1 |  | 18 000 | 0.8 | 1.4 |  |  |  |
| 50.4 |  | 5 925 | 0.6 | 0.3 |  |  |  |
| 35.0 |  | 14 548 | 0.9 | 1.3 |  |  |  |
| 33.3 |  | 4 243 | 0.7 | 0.3 |  |  |  |
| 30.3 |  | 79 337 | 1.0 | 7.6 |  |  |  |
| 28.8 |  | 15 059 | 0.6 | 0.8 |  |  |  |
| 27.8 |  | 15 844 | 0.8 | 1.3 |  |  |  |
| 26.9 |  | 10 417 | 0.9 | 0.9 |  |  |  |
| 25.9 |  | 14 548 | 0.8 | 1.1 |  |  |  |
| 14.8 |  | 3 021 | 0.7 | 0.2 |  |  |  |

**^a^**	The amount of molecule 1 in GalA/GlcA/Rha is calculated by averaging the resolved C1 and C4 peaks.

**^b^**	The amount of Rha is calculated by averaging the resolved C1 and C6 (CH_3_) peaks.

**^c^**	In method 1, the amount of cellulose is calculated by adding all the C4 resonances (interior and surface chains).

**^d^**	In method 2, the amount of cellulose is calculated using the relatively resolved i6 peak, and the back calculate the amount of all cellulose using the interior-to-surface ratio obtained in CP spectrum.

**^e^**	Galactose is component added in the last step, with high uncertainty. Therefore, intensities of all carbon sites are averaged.

**^f^**	The amount of arabinose is calculated by averaging the C1, C2 and C4 peaks.

**^g^** The backbone of xyloglucan (XyG) is the average of the C1 and C4 peaks.

**Additional file 4. Table S3.** Molecular composition derived from ^13^C MultiCP spectrum in Figure 3b, c. The relatively well resolved resonances (underlined) are used for estimating polysaccharide composition. Carbohydrate peaks account for 85% of total intensity and lipid polymers account for 15% of all carbons.

|  | Carbohydrate | Lipid |
| --- | --- | --- |
| Fraction | 84.5% | 15.5% **^a^** |

|  | GalA/GlcA | Rha | Cellulose **^b^** | Gal | Ara | XyG **^c^** |
| --- | --- | --- | --- | --- | --- | --- |
| Method 1 | 47% | 3% | 28% | 9% | 6% | 6% |
| Method 2 | 42% | 3% | 35% | 8% | 5% | 6% |

**^a^**	The carbohydrate resonances in **Additional file 3** account for 84.5% of all carbons. The lipid resonances account for 15.5 % of all carbons in the sample.

**^b^**	In Method 1, the cellulose content is calculated by adding all the underlying carbon 4 resonances from 84 to 90 ppm. In Method 2, the cellulose content is calculated using only the signal from i6 peak in MultiCP, and the interior-to-surface ratio in CP.

**^c^**	XyG amount is the sum of xylose and glucose in **Additional file 3**. Note that significant uncertainty is present with the amount of XyG as it is poorly resolved.

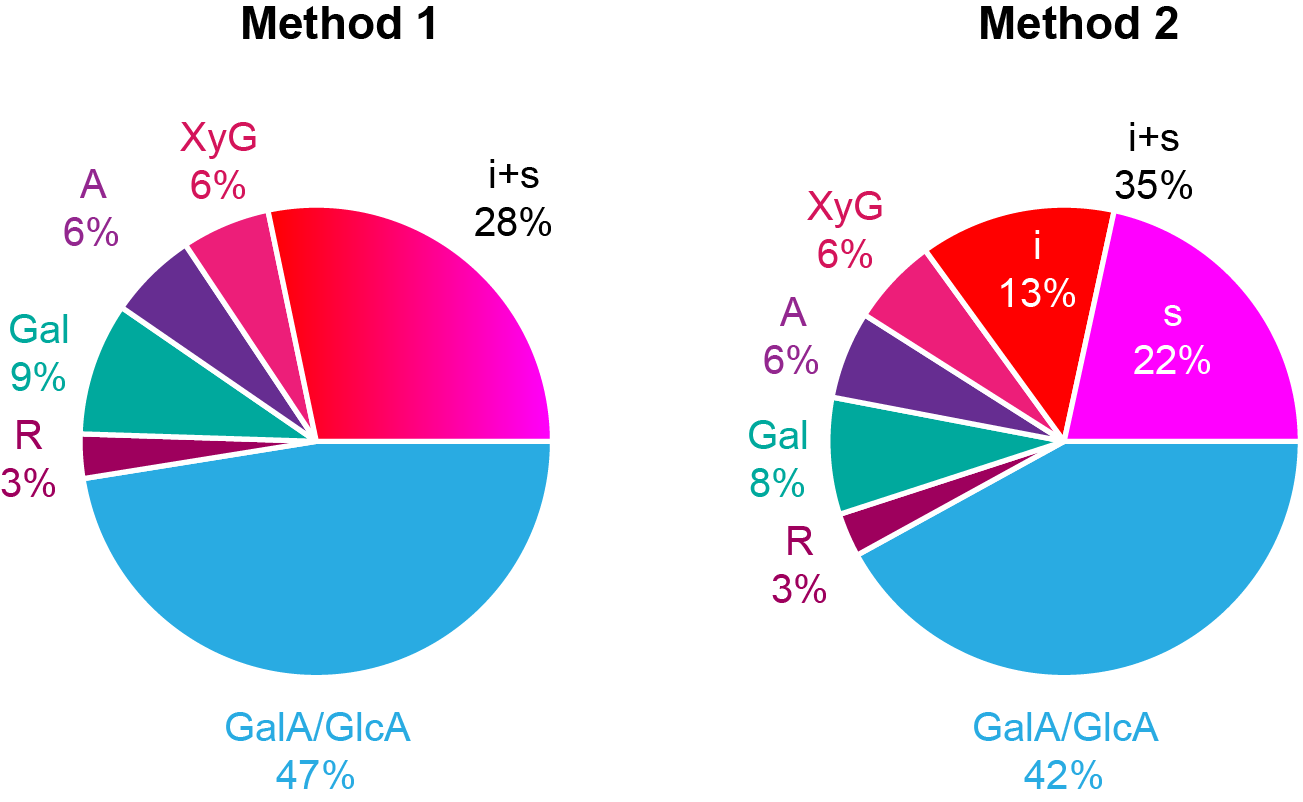

**Additional file 5. Figure S1.** MultiCP integration models. **Method 1:** For each sugar, except cellulose, percentages are obtained by averaging MultiCP integrals of resolved peaks (underlined values in **Additional file 3**, to obtain each sugar contribution), and dividing the result by the sum of all contributions (including cellulose one). For cellulose, as C4 region of MultiCP is broad and poorly defined, i and s contributions come from the sum of all peaks in this region. **Method 2:** Sugar contributions are obtained with the same method. However, for cellulose, this time resolved s6 peaks on MultiCP is used as a reference. i and s contributions are obtained by multiplying this integral by i to s ratio obtained from the usual ratio of C4 integrals in the classic CP experiment. We note that this second method clearly over-estimate total cellulose content as i6 MultiCP deconvolution component is particularly difficult to obtain. i (interior cellulose), s (surface cellulose), Gal (galactose), A (arabinose), R (rhamnose), and XyG (xyloglucan).
